# Multiplexed and Inducible Gene Modulation in Human Pluripotent Stem Cells by CRISPR Interference and Activation

**DOI:** 10.1101/603951

**Authors:** Dane Z. Hazelbaker, Amanda Beccard, Patrizia Mazzucato, Gabriella Angelini, Angelica Messana, Daisy Lam, Kevin Eggan, Lindy E. Barrett

## Abstract

CRISPR-Cas9-mediated gene interference (CRISPRi) and activation (CRISPRa) approaches hold promise for functional genomic studies and genome-wide screens in human pluripotent stem cells (hPSCs). However, in contrast to CRISPR-Cas9 nuclease approaches, the efficiency of CRISPRi/a depends on continued expression of the dead Cas9 (dCas9) effector and guide RNA (gRNA), which can vary substantially depending on transgene design and delivery. Here, we design new fluorescently labeled *piggyBac* (PB) vectors to deliver robust and stable expression of multiplexed gRNAs. In addition, we generate hPSC lines harboring AAVS1-integrated, inducible and fluorescent dCas9-KRAB and dCas9-VPR transgenes to allow for accurate quantification and tracking of cells that express both the dCas9 effectors and gRNAs. We then employ these systems to target the *TCF4* gene and conduct a rigorous assessment of expression levels of the dCas9 effectors, gRNAs and targeted gene. Collectively, these data provide proof-of-principle application of a stable, multiplexed PB gRNA delivery system that can be widely exploited to further enable genome engineering studies in hPSCs. Paired with diverse CRISPR tools including our dual fluorescence CRISPRi/a cell lines, this system would facilitate functional dissection of individual genes and pathways as well as larger-scale screens for studies of development and disease.

## INTRODUCTION

CRISPR-Cas9 systems have revolutionized genome editing in myriad cell types and organisms and ushered the development of variant technologies that utilize dCas9 fused to epigenetic modifiers which can be localized to a gene of interest upon expression of a gRNA (Adli, 2018; Chavez et al., 2015; Gilbert et al., 2014). Two such approaches are CRISPRi, which fuses dCas9 to transcriptional repressors, such as the KRAB domain (Gilbert et al., 2014), and CRISPRa which fuses dCas9 to transcriptional activators, such as the chimeric VPR domain (Chavez et al., 2015). These tools can be deployed for both single and multiplexed gene manipulation and allow modulation of gene expression in the absence of cellular toxicity caused by Cas9-mediated DNA double-strand breaks (Aguirre et al., 2016). CRISPRi/a set-ups have been used successfully in studies of cellular programming (Kearns et al., 2013), cellular reprogramming (Liu et al., 2018; Weltner et al., 2018), *in vivo* gene manipulation (Zhou et al., 2018), enhancer screens (Fulco et al., 2016), chemical screens (Jost et al., 2017), and whole-genome genetic interaction mapping studies (Horlbeck et al., 2018). When targeting populations of cells, gene repression through CRISPRi is reported to be more homogeneous and efficient compared to Cas9 nuclease (Mandegar et al., 2016). Indeed, while Cas9-nuclease strategies have been employed in genome-wide screens, they are limited by heterogeneity in the targeted cell populations, which may include a significant number of wild-type cells alongside cells with mixtures of indels that produce partial loss or gain of function phenotypes, or truncated gene products which can complicate interpretations (Mandegar et al., 2016). Furthermore, CRISPRi/a offers the potential for conditional gene perturbation, allowing for the functional study of essential genes (Gilbert et al., 2014) and reversibility of phenotypes. However, unlike genetic knockout by CRISPR-Cas9 that requires a single indel formation event to permanently disrupt gene function, successful CRISPRi/a requires persistent and uniform expression of dCas9 effectors and gRNA across cell populations, an important consideration both in single gene studies and whole-genome screens.

There is limited data on the stability of dCas9 effectors (Mandegar et al., 2016) and studies report variability in the induction and expression of different promoters in different loci due to *de novo* DNA methylation (Bertero et al., 2016a). Further, gRNA delivery and expression require optimization in order to fully capitalize on the multiplexing potential of CRISPRi/a. With regard to gRNA delivery, previous studies have utilized transfection and selection of plasmid DNA (Balboa et al., 2015; Heman-Ackah et al., 2016; Mandegar et al., 2016) transient transfection of *in vitro* transcribed gRNA (González et al., 2014; Ho et al., 2017), lentiviral integration (Ho et al., 2017) or *piggyBac* transposon-based integration (Li et al., 2017). In particular, *piggyBac* (PB) delivery methods have the advantages of being easy to clone and deliver into hPSCs and carry substantially larger payload compared to lentiviral vectors (Schertzer et al., 2018; Wang et al., 2016). As a result, PB vectors are particularly applicable for studies of parallel pathways or polygenic disease, enabling the perturbation of many genes with a single delivery vehicle at minimal cost.

Here, we developed a new *piggyBac* vector system to enable rapid cloning and stable delivery of multiple gRNAs for CRISPRi/a applications. We coupled this system with genomically integrated and inducible dCas9-KRAB and dCas9-VPR in hPSCs, including a dual-fluorescent readout to readily quantify cells that express both gRNAs and dCas9 variants in a population. We then quantified expression levels of the effector components as well as a targeted gene, *TCF4*, at both the transcript and protein levels. Our results confirm the utility of the dual-fluorescent readout and multiplexed PB gRNA delivery system for CRISPRi/a that can now be broadly employed in hPSCs for gene perturbation studies.

## RESULTS

### Generation of AAVS1 integrated and doxycycline-inducible dCas9-KRAB and dCas9-VPR hPSC lines

To derive stable CRISPRi and CRISPRa hPSC lines, we cloned and introduced all-in-one cassettes containing S. *pyogenes* dCas9 fused to the KRAB repressor domain (Gilbert et al., 2013) or VPR activation domain (Chavez et al., 2015) into the AAVS1 safe-harbor locus of the XY embryonic stem cell line H1 (Thomson et al., 1998) via a TALEN-mediated gene-trap approach that confers neomycin (G418) resistance to cells upon on-target integration (González et al., 2014; Mandegar et al., 2016) (**Figure 1A**). In both constructs, dCas9-KRAB and dCas9-VPR expression is driven by the TRE3G doxycycline inducible promoter (Takara Bio) and fused to Enhanced Green Fluorescent Protein (EGFP) transcriptional reporters by an IRES sequence (dCas9-KRAB) or a T2A self-cleaving peptide sequence (dCas9-VPR). Following selection with G418, dCas9-KRAB and dCas9-VPR clones were assessed for EGFP expression and genotyped by junction PCR (**Figure S1A, B**). From these data, dCas9-KRAB and dCas9-VPR clones were expanded and confirmed to have normal karyotypes and absence of mycoplasma (data not shown).

**Figure 1.**
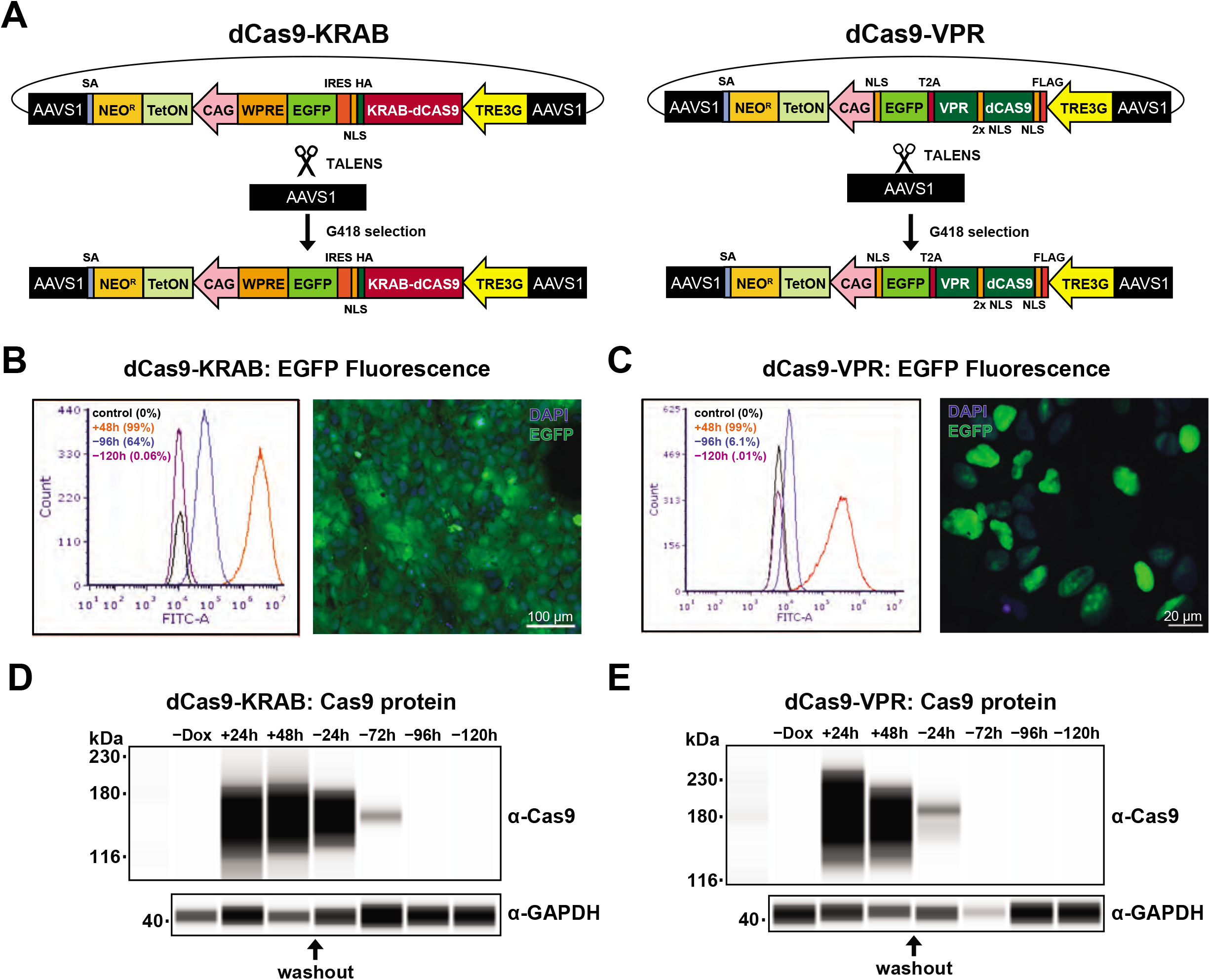
Generation and validation of AAVS1-integrated inducible dCas9-KRAB and dCas9-VPR systems in hPSCs. **A.** Schematic overview of AAVS1 targeting strategy in H1 hPSCs with TRE3G-driven dCas9-KRAB (left) or dCas9-VPR (right) cassettes and TALENs that target AAVS1 and confer G418 resistance upon on-target integration. **B.** *Left*, Flow cytometry analysis of EGFP fluorescence in dCas9-KRAB cells after 48 hours of doxycycline treatment (+48h) followed by removal of doxycycline for 96 (−96h) and 120 hours (−120h) in comparison to no GFP control H1 cells (control). *Right*, representative image of EGFP expression in dCas9-KRAB cells after 48 hours doxycycline treatment. **C.** *Left*, Flow cytometry analysis of EGFP fluorescence in dCas9-VPR cells after 48 hours of doxycycline treatment (+48h) followed by removal of doxycycline for 96 (−96h) and 120 hours (−120h) in comparison to no GFP control H1 cells (control). *Right*, representative image of EGFP expression in dCas9-VPR cells after 48 hours doxycycline treatment. **D.** dCas9-KRAB protein expression in absence of doxycycline (-Dox), after 24 and 48 hours doxycycline treatment (+24h, +48h), and after washout of doxycycline for 24, 72, 96, and 120 hours (−24h, −72h, −96h, −120h). **E.** dCas9-VPR protein expression before doxycycline treatment (-Dox), after 24 and 48 hours doxycycline treatment (+24h, +48h) and after washout of doxycycline for 24, 72, 96, and 120 hours (−24h, −72h, −96h, −120h).

To validate our CRISPRi/a hPSC lines, we first quantified EGFP fluorescence by flow cytometry following 48 hours of doxycycline treatment. Doxycycline led to strong induction of EGFP fluorescence, reaching 99% in both dCas9-VPR and dCas9-KRAB hPSC lines (+48h; **Figure 1B, C**). 120 hours after washout of doxycycline, EFGP fluorescence levels dropped to background levels in both the dCas9-KRAB and dCas9-VPR lines (−120h; **Figure 1B, C**). As expected, we also observed strong induction of dCas9-KRAB and dCas9-VPR protein expression after doxycycline induction (**Figure 1D, E**) and loss of detectable dCas9 expression by 96 hours post-washout in dCas9-KRAB cells and by 72 hours in dCas9-VPR cells (**Figure 1D, E**). The increased stability of dCas9-KRAB and EGFP protein in dCas9-KRAB cells in comparison to dCas9-VPR cells may be due to the presence of the WPRE (<u>W</u>oodchuck Hepatitis Virus <u>P</u>ost-transcriptional <u>R</u>esponse <u>E</u>lement) in the 3’ UTR of the dCas9-KRAB construct (**Figure 1A**), which has been reported to increase transcript stability (Zufferey et al., 1999). Similar to previous reports, dCas9 protein was not detected in the absence of doxycycline (Mandegar et al., 2016)(**Figure 1D, E**). These data confirm that our AAVS1-integrated dCas9-KRAB and dCas9-VPR constructs exhibit robust induction and reversibility of dCas9 expression in hPSCs.

### Identification of the relevant transcriptional start site of *TCF4* in hPSCs

To assess the potency of our dCas9-KRAB and dCas9-VPR systems for gene repression and activation, we targeted the *TCF4* gene in hPSCs as an example. *TCF4* plays important roles in development and *TCF4* gene dysfunction has been implicated in multiple neurodevelopmental diseases including Pitt-Hopkins syndrome and schizophrenia by GWAS (Jung et al., 2018; Quednow et al., 2014; Ripke et al., 2014). Importantly, *TCF4* has multiple alternatively-spliced transcripts (Sepp et al., 2011) making it critical to identify the most relevant TCF4 isoform and its corresponding transcriptional start site (TSS) to target with CRISPRi/a. Generally speaking, functional gRNA design for CRISPRi/a applications has the added challenge that TSSs may not be well annotated for a given cell type. We therefore first carried out western blot analysis of TCF4 protein in hPSC lysates. As shown in **Figure 2A**, the most abundant and full-length isoform migrated at approximately 72 kDa, which corresponds to the full-length canonical TCF4 sequence (Sepp et al., 2011). To experimentally map the functional TSS of this protein isoform, we utilized exon-specific RT-qPCR with an array of primers targeting candidate TSS-harboring exons. RT-qPCR analysis revealed the most dominantly expressed exon to be exon 3b of *TCF4*, which corresponds to the TCF4-B transcript isoform (**Figure 2B**) (Sepp et al., 2011). Having established the dominant TSS of *TCF4* in hPSCs, we selected three gRNAs for CRISPRi (i1, i2, i3) and three gRNAs for CRISPRa (a1, a2, and a3) that fall within the optimal window of 300 bp from the TSS (Doench, 2017; Gilbert et al., 2014) using the CRISPR-ERA guide selection tool (Liu et al., 2015)(**Figure S2A**).

**Figure 2.**
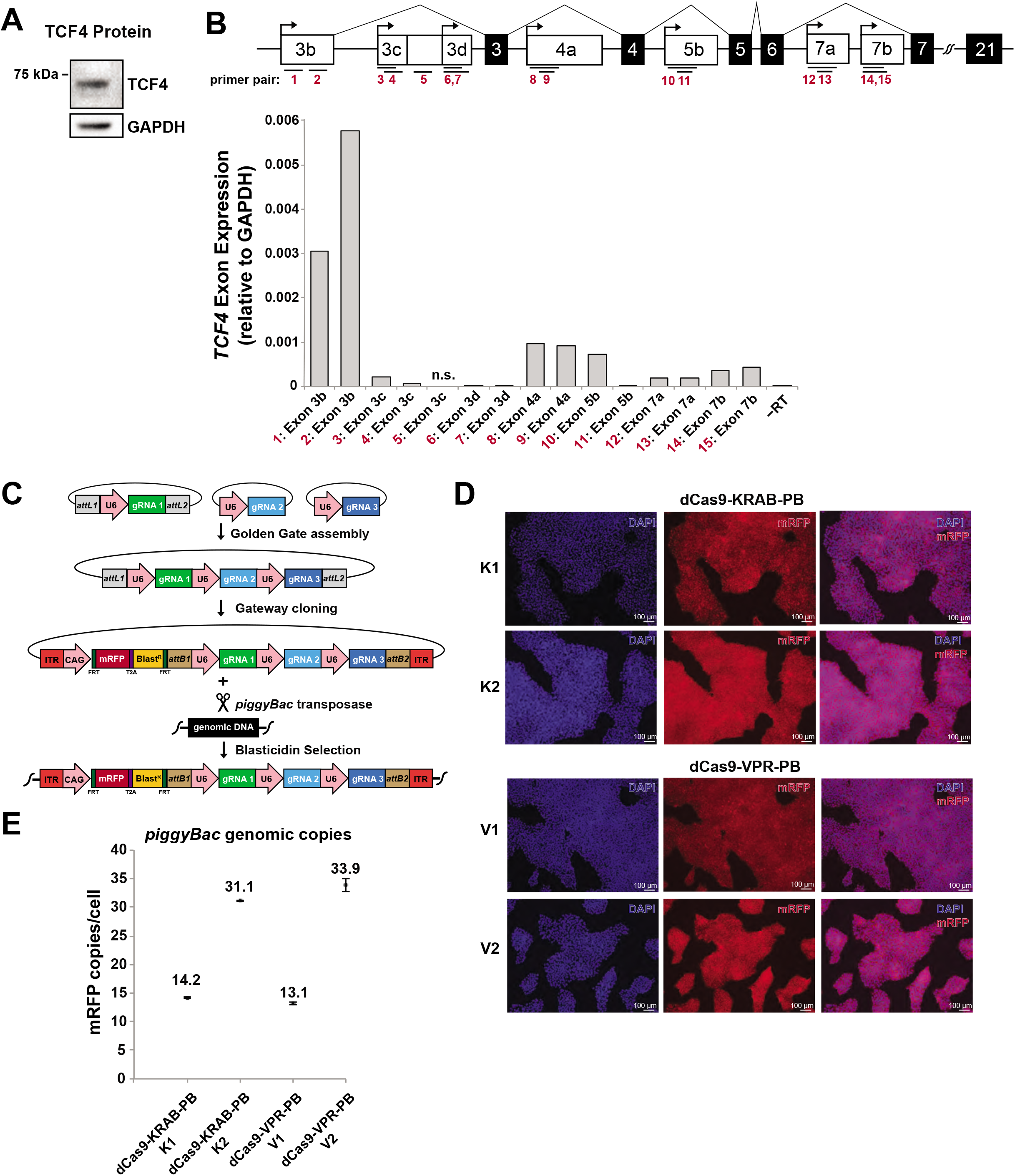
Design and delivery of multi-gRNA PB vectors for CRISPRi and CRISPRa targeting of the *TCF4* gene. **A.** Representative western blot of TCF4 protein expression in hPSCs. **B.** *Top*, Schematic of the *TCF4* gene with primer pairs in red corresponding to exon locations modified from (Sepp et al., 2011). *Bottom*, Exon-specific expression of *TCF4* transcripts in hPSCs compiled from RT-qPCR of H1 cells and normalized to GAPDH. n.s. corresponds to no signal in the qPCR reaction. **C.** Overview of multi-gRNA PB vector cloning, delivery, and selection. **D.** Representative images of mRFP fluorescence in dCas9-KRAB-PB clones K1 and K2 (*top panels*) and dCas9-VPR-PB clones V1 and V2 (*bottom panels*). Cells are counterstained with DAPI (blue). Scale bar = 100μm. **E.** PB vector copy number in dCas9-KRAB and dCas9-VPR clones as determined by ddPCR quantification of mRFP gene. Data is shown as the mean of three experiments with error bars as +/− s.e.m.

### Design and delivery of multi-gRNA *piggyBac* vectors in hPSCs

A noted strength of CRISPRi and CRISPRa is the ability to deliver multiple gRNAs for enhanced targeting of one or several genes in the absence of DNA damage (Jusiak et al., 2016). To facilitate stable delivery of multiple gRNAs in hPSCs, we designed a new vector that incorporates the efficiency and ease of the *piggyBac* (PB) transposase system (Chen et al., 2010) with a multiplex gRNA cloning system (Sakuma et al., 2014). To do so, we cloned either the three CRISPRi gRNAs (i1, i2, i3) or CRISPRa gRNAs (a1, a2, a3) targeting *TCF4* into individual vectors and sequentially assembled the final PB vector including mRFP and blasticidin resistance via Golden Gate and Gateway cloning (**Figure 2C**). Of note, while we opted to introduce three gRNAs per PB vector, the parental vectors allow for the cloning of up seven gRNAs in tandem array (Sakuma et al., 2014) that can easily be introduced into our PB vectors. Additionally, *FRT* sites flanking the mRFP and blasticidin cassettes (**Figure 2C**) allow for removal of these selectable features by introduction of FLP recombinase, allowing for future PB re-targeting events. With our workflow, the cloning of multiple gRNAs into these vectors can be completed and confirmed via BamHI restriction digest within one week (**Figure S2B and Materials and Methods**).

We next co-transfected the multi-gRNA PB vectors along with a plasmid encoding the *piggyBac* transposase into dCas9-KRAB and dCas9-VPR hPSC lines. Following selection with blasticidin, individual dCas9-KRAB and dCas9-VPR clones were isolated and screened for high levels of uniform mRFP fluorescence (**Figure 2D**). We then selected and expanded two independent dCas9-KRAB clones (dCas9-KRAB-PB clones K1 and K2) and two independent dCas9-VPR clones (dCas9-VPR-PB clones V1 and V2) and assessed integrated PB copy number via droplet digital PCR (ddPCR). Our ddPCR analysis revealed approximately 14 and 31 PB copies in dCas9-KRAB-PB clones K1 and K2, respectively, and approximately 13 and 34 PB copies dCas9-VPR-PB clones V1 and V2, respectively (**Figure 2E**). At this stage, we also confirmed that both dCas9-KRAB-PB and dCas9-VPR-PB cells harboring moderate levels of integrated PBs (i.e., 13 - 34 copies) maintained pluripotency and tri-lineage potential (**Figure S2C, D**). Thus, our multiplexed PB vectors can facilitate rapid cloning and efficient delivery of gRNAs for CRISPRi/a applications in hPSCs.

### Quantifying CRISPRi/a Component Expression in hPSCs

To quantify CRISPRi/a component expression, that is, the dCas9 effector and gRNA levels, we treated dCas9-KRAB-PB and dCas9-VPR-PB clones with doxycycline for 0, 24, and 48 hours and collected replicate and matched samples for side-by-side clonal analysis via flow cytometry, western blot, and RT-qPCR (**Figure 3A**). For CRISPRi, flow cytometric quantification of EGFP fluorescence of the two independent dCas9-KRAB-PB clones showed high levels of EGFP fluorescence (99.9% for both K1 and K2 clones) and mRFP expression fluorescence (100% for both K1 and K2 clones) after 48 hours of doxycycline induction (**Figure 3B**) indicating robust and uniform expression of dCas9-KRAB and gRNA. In direct congruence with the EGFP and mRFP fluorescence data, we observe strong induction of dCas9-KRAB protein in both K1 and K2 clones (**Figure 3C**) and high levels of all three CRISPRi gRNAs by gRNA-specific RT-qPCR (**Figure 3D**). Indeed, we observed gRNA expression levels between 1% and nearly 100% of the levels of *GAPDH* transcripts. These results indicate that PB vectors provide a consistent and reproducible means to express multiple gRNAs across cells in a population using a single delivery vehicle.

**Figure 3.**
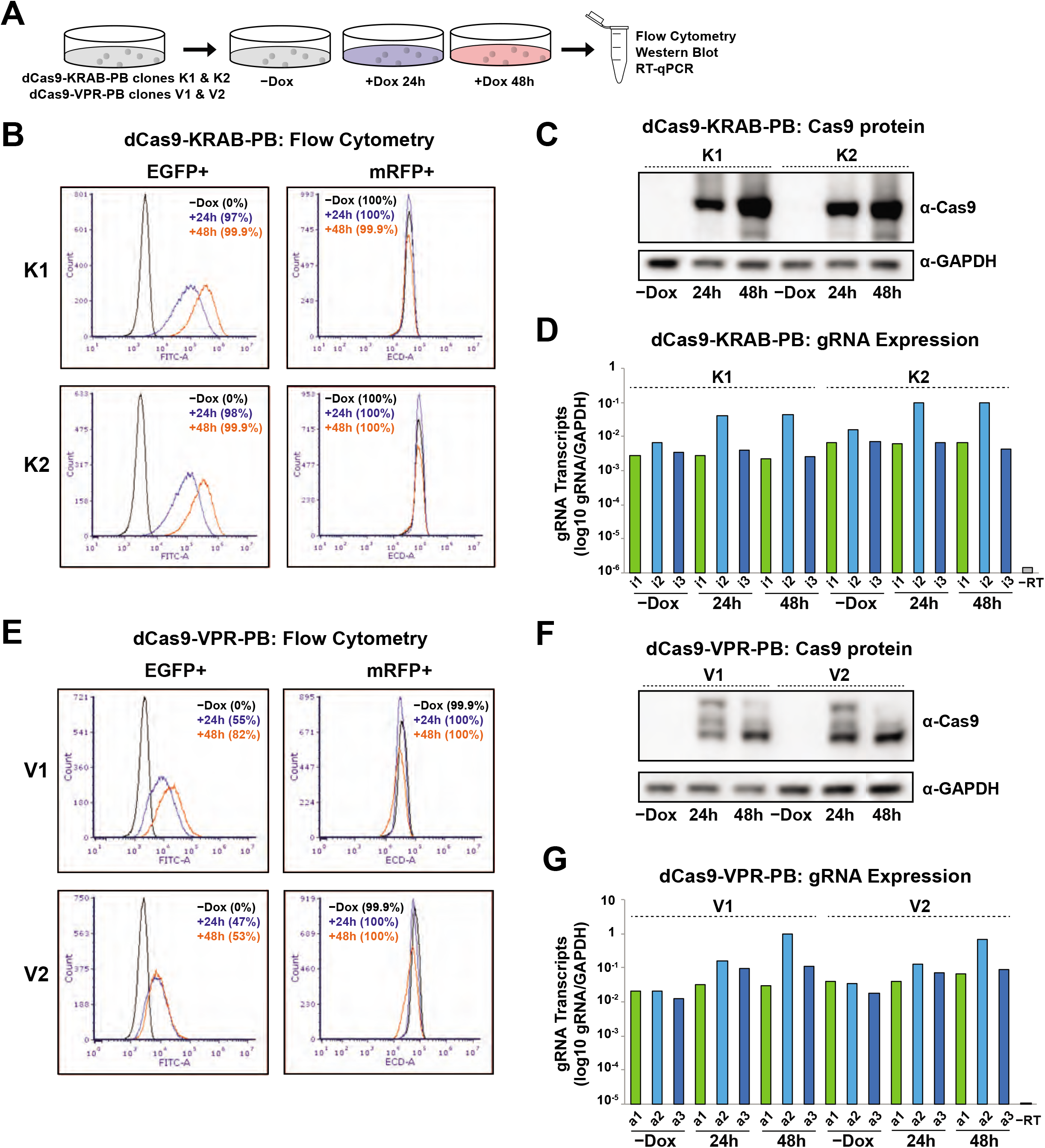
Assessment of CRISPRi and CRISPRa component expression. **A.** Experimental overview for activation and repression of *TCF4* in dCas9-KRAB-PB and dCas9-VPR-PB clones **B.** Flow cytometry analysis of EGFP and mRFP fluorescence in dCas9-KRAB-PB clones in absence of doxycycline (-Dox) and in presence of doxycycline for 24 (+24h) and 48 (+48h) hours. **C.** Western blot analysis of dCas9-KRAB protein in dCas9-KRAB-PB clones K1 and K2 at indicated time-points. **D.** RT-qPCR analysis of gRNAs i1, i2, and i3 in dCas9-KRAB-PB clones K1 and K1. **E.** Flow cytometry analysis of EGFP and mRFP fluorescence in dCas9-VPR-PB clones at indicated time-points. **F.** Western blot analysis of dCas9-VPR protein level in dCas9-VPR-PB clones at indicated time-points. **G.** RT-qPCR analysis of gRNAs a1, a2, and a3 expression in dCas9-VPR-PB clones V1 and V2.

In the case of CRISPRa, flow cytometric quantification of EGFP fluorescence of the two independent dCas9-VPR-PB clones revealed only intermediate levels of EGFP fluorescence (82% for clone V1 and 53% for clones V2; **Figure 3E**). The decreased levels of EGFP expression in dCas9-VPR-PB clones contrasts with the high levels of EGFP expression in the parental dCas9-VPR clone (**Figure 1C**), perhaps indicating a CRISPRa gRNA- or PB-specific effect as prolonged doxycycline treatment in dCas9-VPR cells without integrated PB vectors did not result in reduced EGFP levels (data not shown). However, mRFP expression remained high in dCas9-VPR-PB cells, as quantified by flow cytometry (100% for both clone V1 and V2). Direct assessment of dCas9-VPR protein levels in clones V1 and V2 revealed strong induction upon doxycycline treatment (**Figure 3F**), albeit to a lesser extent compared with dCas9-KRAB-PB clones. Again, RT-qPCR confirmed robust and uniform expression of all three CRISPRa gRNAs (**Figure 3G**). Interestingly, the expression levels of CRISPRi and CRISPRa gRNAs i2 and a2 increased 4-6 fold upon expression of dCas9 by doxycycline treatment (**Figures 3D, G**). It is possible that the presence of dCas9 selectively increases gRNA stability by binding particular gRNAs with high affinity and protecting them from degradation, perhaps by masking the 5’ end of the gRNA, as suggested by previous studies (Jiang and Doudna, 2017). These results demonstrate that both dCas9-effectors and multiplex gRNAs are efficiently expressed in our CRISPRi and CRISPRa hPSC lines.

### Quantification of TCF4 repression and activation at the transcript and protein level

Having established robust expression of our effector components, we next sought to quantify levels of repression and activation of a target gene in hPSCs. To quantify the efficiency of *TCF4* repression by CRISPRi, we first analyzed *TCF4* transcript levels by RT-qPCR in dCas9-KRAB-PB clonal pairs. As shown in **Figure 4A**, doxycycline induction resulted in rapid and significant repression of TCF4 transcripts both clones K1 and K2, (averaged decreased of 18-fold at 24 hours and 200-fold at 48 hours). Comparison of TCF4 transcripts in clone K1 and K2 shows that K2 displays more rapid repression, suggesting that CRISPRi potency may titrate with PB copy number (**Figure 2E**). In contrast to the rapid decline of *TCF4* transcripts, TCF4 protein was more moderately decreased in both dCas9-KRAB clones, resulting in a reduction of 0.4-fold after 24 hours and 2-fold after 48 hours of doxycycline treatment (**Figure 4B**). By comparison, targeting of *TCF4* for activation in dCas9-VPR-PB clones resulted in a 1.8- and 1.6-fold averaged increase in transcript levels after 24 hours and 48 hours doxycycline treatment, respectively (**Figure 4C**). TCF4 protein levels increased approximately 2- and 1.3-fold after 24 and 48 hours of doxycycline induction, respectively, in dCas9-VPR-PB cells (**Figure 4D**).

**Figure 4.**
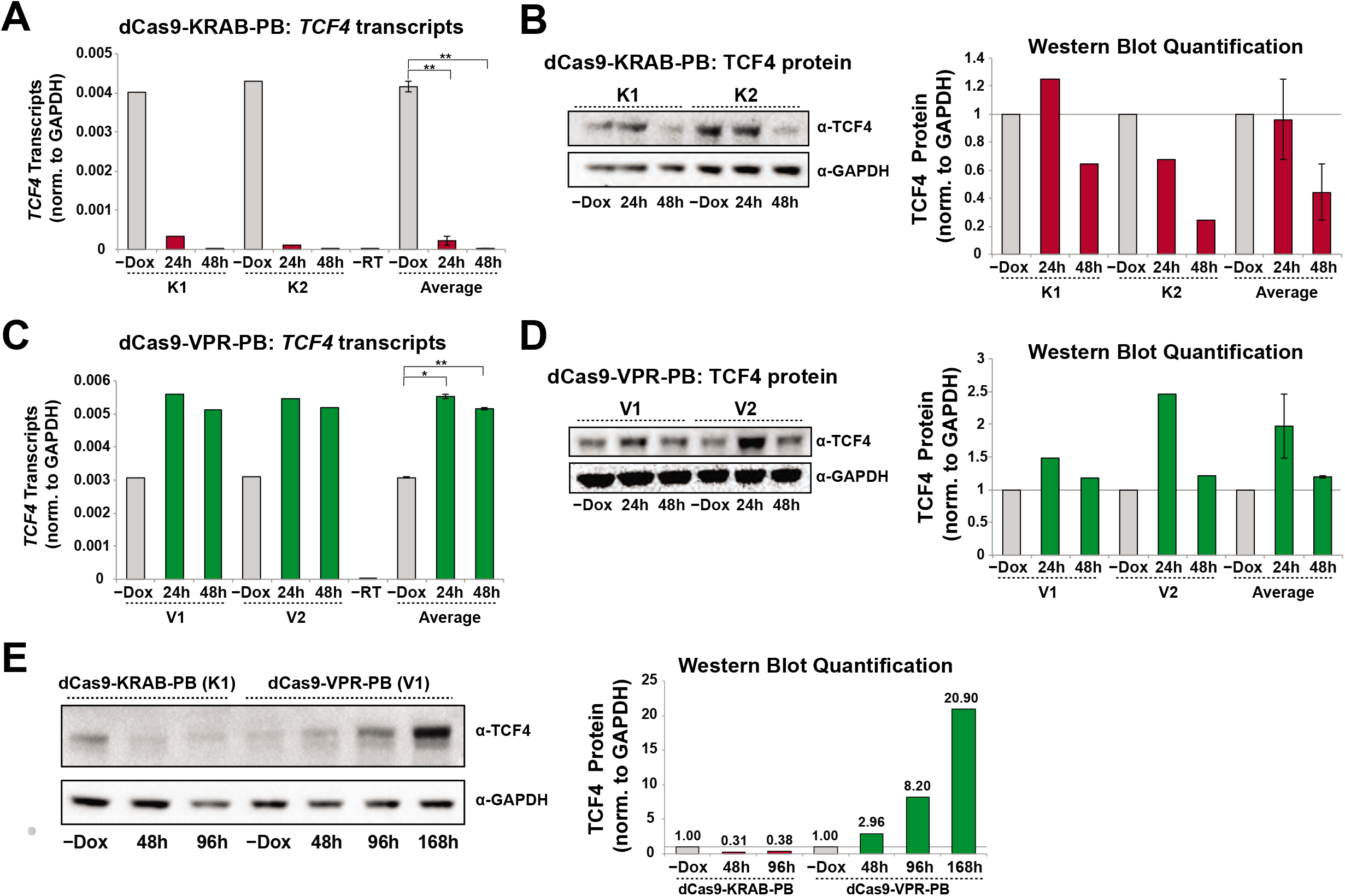
Repression and activation of *TCF4* at the transcript and protein levels. **A.** RT-qPCR analysis of *TCF4* transcript levels at indicated time points in dCas9-KRAB-PB clones K1 and K2 separately and averaged. Data is shown as mean +/− s.e.m. **B.** Western blot analysis of TCF4 protein in dCas9-KRAB-PB clones K1 and K2 at indicated time-points with quantification shown on the right. **C.** RT-qPCR analysis of *TCF4* transcript levels at indicated time points in dCas9-VPR-PB clones V1 and V2 separately and averaged. Data is shown as mean +/− s.e.m. **D.** Western blot analysis of TCF4 protein in dCas9-VPR-PB clones V1 and V2 at indicated time-points with quantification shown on the right. **E.** Western blot analysis of TCF4 protein levels in extended course of doxycycline treatment in dCas9-KRAB-PB clone K1 and dCas9-VPR-PB clone V1 at indicated time-points with quantification shown on the right. In all panels, expression levels are normalized to GAPDH and *P < 0.05, **P < 0.01 (two tailed paired T test).

To confirm our results over longer induction time-points, we treated the dCas9-KRAB-PB clone K1 with doxycycline for up to 96h and the dCas9-VPR clone V1 for up to 168h. As shown in **Figure 4E**, the 96h induction period for dCas9-KRAB-PB cells failed to further decrease TCF4 protein levels (~3-fold reduction at 48h *versus* 2.5-fold reduction after 96h). By contrast, continued doxycycline induction of dCas9-VPR resulted in a steady increase of TCF4 protein over time, with a 20-fold increase after 168h (**Figure 4E**). These results are consistent with activation of *TCF4* through CRISPRa resulting in parallel activation of *TCF4* transcript and protein expression levels, and repression of *TCF4* through CRISPRi leading to differing degrees of impact on transcript and protein expression. These differences likely reflect endogenous gene regulatory programs that remain notable considerations for CRISPRi/a applications. Importantly, our data confirm robust and efficient transcript repression and activation of the target gene, thus providing proof-of-principle data on the effectiveness of our CRISPRi/a approach in hPSCs.

## DISCUSSION

CRISPRi/a systems hold great potential for exploring gene function and dissecting human disease mechanisms in hPSCs and hPSC-derived cell types, such as cardiomyocytes (Mandegar et al., 2016) and neurons (Ho et al., 2017). Benefits of CRISPRi/a over knockout strategies utilizing Cas9 nuclease include the ability to conditionally perturb essential and multiple genes in the absence of DNA damage and genetic instability (Ihry et al., 2018; Kosicki et al., 2018). However, in contrast to gene perturbation by gene knockout with CRISPR-Cas9, gene modulation by CRISPRi and CRISPRa approaches are dependent on sustained expression of the dCas9 effector and gRNA.

Here, we developed a set of tools to facilitate multiplexed CRISPR-mediated gene modulation in hPSCs. We find that our integrated dCas9-KRAB and dCas9-VPR constructs allow for reproducible and reversible induction of dCas9 alongside EGFP in the vast majority of cells, consistent with previous reports (Mandegar et al., 2016). To facilitate stable and multiplex gRNA expression, we designed and validated a drug-selectable *piggyBac* vector with constitutive mRFP fluorescence to visualize and track gRNA-expressing cells. Thus, dual monitoring of both EGFP and mRFP fluorescence allows for quantification of the percentage of CRISPRi/a competent cells in a population. This may be particularly important for downstream functional studies and screens in differentiated cell types derived from hPSCs, where cell-to-cell variation is likely to increase. Further, some commonly used mammalian promoters are reported to be silenced over time by DNA methylation (Bertero et al., 2016b; Norrman et al., 2010) and lentiviral vectors, commonly used to introduce gRNAs, can also be subject to shutdown (Xia et al., 2007).

With regard to gRNA expression, we find that CAG-promoter driven PB vectors support sustained gRNA and reporter expression in hPSCs. Importantly, we confirmed high expression levels of 3 independent gRNAs in the multiplexed system, demonstrating that PB vectors provide a dependable, rapid and inexpensive delivery vehicle for transgene expression. Specifically, we anticipate these vectors will be useful for rapid and multiplexed expression of gRNAs in hPSCs for perturbation analysis at the single gene and whole genome levels, both in CRISPRi/a contexts, as presented here, and in CRISPR knockout schemes with Cas9 nuclease or Cas9-fused base editors (Billon et al., 2017).

In our examination of CRISPRa potency, we observed a near 1-to-1 congruency between level of transcriptional activation and protein overexpression. However, in our assessment of CRISPRi, we find that *TCF4* transcript levels drop precipitously, approximately 200-fold within 48 hours of dCas9-KRAB induction, while protein levels were reduced merely 2-fold. This discrepancy may arise in cases where only a small percentage of the expressed transcripts are needed to maintain cellular protein levels or cases when the protein is more stable than the transcript (Schwanhäusser et al., 2011). Such transcription-translation discrepancies are mediated by endogenous regulatory programs that vary from cell-type to cell-type (Moritz et al., 2019) and remain an important aspect to consider in both single gene studies and whole genome screens with CRISPRi and CRISPRa strategies.

Collectively, our newly designed multi-gRNA PB vectors are vehicles for robust, sustained gRNA expression in hPSCs. Further, the coupling of these tools with our dual-fluorescence dCas9-KRAB and dCas9-VPR systems facilitates accurate quantification and tracking of CRISPRi/a components across cells in a population. We anticipate these tools will facilitate both single and multiplexed gene perturbation studies and screens in hPSCs and other cell types for functional interrogation of development and disease.

## MATERIALS AND METHODS

### Plasmid construction

To generate the dCas9-KRAB-IRES-EGFP AAVS1 targeting plasmid pT077, parental plasmid pHR-TRE3G-KRAB-dCas9-IRES-GFP (a gift from Jesse Engreitz, Broad Institute) was cloned by Gibson assembly into the backbone fragment of plasmid pGEP116 that contains AAVS1 homology arms and the doxycycline-responsive activator rTTA driven by a constitutive CAG promoter (Sellgren et al., 2019). To generate the dCas9-VPR-T2A-EGFP AAVS1 targeting plasmid (pT076), the dCas9-VPR cassette from plasmid SP-dCas9-VPR (Addgene 63798, (Chavez et al., 2015)) was fused by Gibson assembly with a PCR fragment containing a T2A-EGFP-NLS cassette (from plasmid PT059) and cloned into plasmid pGEP116. Oligonucleotides (IDT) corresponding to gRNA target sequences (Supplemental Table S1) were cloned via BpiI into pX330S-2 and pX330S-3 (Sakuma et al., 2014) and a third vector pGEP179_pX330K (this study) according to kit instructions (Addgene Kit#1000000055, Sakuma et al., 2014). The pGEP179_pX330K plasmid is a modified entry vector generated by cloning the BsaI-pU6-sgRNA-BsaI fragment from pX330A-1×3 (Sakuma et. al., 2014) into a slightly modified MCS of the *attL-containing* entry vector, pENTR1A (Invitrogen). These gRNA-containing pX330S and pGEP179_pX330K plasmids were then assembly by Golden Gate cloning to form a single entry vector. This entry vector was then cloned by Gateway cloning into the *piggyBac* donor destination plasmid pGEP163 that contains *piggyBac* ITRs for transposase-mediated insertion, a CAG promoter driving an mRFP-T2A-BLAST^R^ cassette, and *attR* sites Gateway cloning to create CRISPRi multi-gRNA plasmid pPN441 and CRISPRa multi-gRNA plasmid pPN440. Donor plasmid pGEP163 was constructed by fusing a fragment of plasmid PB-CA (Addgene 20960, (Woltjen et al. 2009) containing the *piggyBac* ITRs and a CAG promoter with a synthetic gene block containing FRT-mRFP-T2A-BLAST^R^-SV40 pA-FRT (IDT). The versions of pPN441 and pPN440 plasmids used in this study were initially created by an earlier cloning strategy that was replaced by the strategy described above to generate the same pPN441 and pPN440 plasmids more rapidly.

### Cell culture and gene targeting

The human embryonic stem cell line H1 (WA01) was obtained from WiCell Research Institute (Madison, WI) (Thomson et al., 1998). Stem cells were grown in either mTeSR1 medium (Stem Cell Technologies 05850) or StemFlex medium (ThermoFisher A3349401) on Geltrex (Life Technologies A1413301) coated plates under conditions previously described (Hazelbaker et al., 2017). Throughout culturing, cells were tested to confirm the absence of mycoplasma contamination (Lonza MycoAlert LT07-418). To integrate the dCas9-KRAB and dCas9-VPR constructs into the AAVS1 locus, 2.5 x 10^6^ cells were co-transfected with 10ug of pT077 (KRAB) or pT076 (VPR), 1.5 μg AAVS1 TALEN L (Addgene 59025) and 1.5 μg AAVS1 TALEN R (Addgene 59026) via the Neon Electroporation System (ThermoFisher) at 1050 mV, 30 ms, 2 pulses. For the first round of clonal selection, the transfected cells were plated at low-density (8,000 cells in a 10cm dish) under G418 selection (50ug/ml, Gibco 10131035) to allow for single-cell colony formation (~10 days). Importantly, cells with the dCas9-KRAB and dCas9-VPR cassettes are kept under selection with G418 for the duration of culture and experiments to protect against shutdown of the AAVS1 integrated transgenes. In this strategy, colonies are picked and deposited into a 96-well plate and when sufficiently dense, the 96-well plate is triplicated to create 3 plates of identical clones. Plate 1 is frozen for storage, plate 2 is treated with doxycycline (Sigma, D9891-25g) at a final concentration of 2 μg/ml 24 hours after duplication for visualization of EGFP+ colonies (with high levels of EGFP expression serving as a proxy for high dCas9 expression), and plate 3 is maintained for expansion and banking of EGFP+ colonies (n=6) while the analysis of plate 2 is performed. For integration of the multiplex PB vectors, 2.5 x 10^6^ dCas9-KRAB and dCas9-VPR cells were transfected with 5 μg of pPN441 (CRISPRi multi-gRNA plasmid) and 5 μg of pPN440 (CRISPRa multi-gRNA plasmid), respectively, with 1 μg of transposase plasmid (System Biosciences #PB210PA-1) under conditions described above. 24 hours after transfection, cells are treated with blasticidin at a final concentration of 2 μg/ml for 12-15 days to select for positive *piggyBac* integrants and allow clearing of free plasmid. Genomic DNA for PCR-based genotyping and *piggyBac* copy number analysis by ddPCR was isolated via the DNeasy Blood and Tissue Kit (Qiagen 69504). For doxycycline induction of dCas9-KRAB and dCas9-VPR, cells are treated with 2 μg/ml doxycycline and pelleted at indicated time points.

### Western blot analysis

To isolate protein for western blot analysis, hPSCs were lysed using Pierce IP lysis buffer (Life Technologies 87787) with protease inhibitors (Sigma Aldrich 11836153001). 20 μg total protein, as determined by Pierce BCA Protein Assay kit (Thermo Scientific 23227), was loaded onto Bolt 4-12% NuPAGE Bis-Tris Plus gels (Invitrogen). Gels were transferred overnight at 4°C to nitrocellulose membranes in 1X NuPAGE transfer buffer (Invitrogen) with 10% methanol. The following antibodies were used for western blot analysis: Cas9 (Diagenode C15310258, 1:1000) TCF4 (Abcam ab217668, 1:500), GAPDH (EMD MAB374; 1:2000), α-rabbit HRP-linked F(ab’)2 (GE Life Sciences NA9340; 1:5000) and α-mouse HRP-linked F(ab’)2 (GE Life Sciences NA9310; 1:5000). Blots were visualized by chemiluminescence with the SuperSignal West Femto kit (Pierce) and imaged and quantified with a ChemiDoc MP Imaging System (BioRad). For quantification of Cas9 protein in dCas9-KRAB and dCas9-VPR parental clones, 1.2 μg total protein was analysed with Cas9 (Diagenode C15310258, 1:400) and GAPDH (EMD MAB374; 1:50) antibodies using the Wes capillary immunoassay system (ProteinSimple)

### Flow cytometry

Flow cytometry was performed at the Broad Institute Flow Facility on a CytoFLEX flow cytometer (Beckman Coulter). Cells were treated with 10 mM ROCK inhibitor (Y-27632) for 4 to 6 hours prior to flow. For each experiment, 100,000 events were recorded and analyzed with FCS Express 6 software (De Novo Software)

### Genomic DNA isolation and genotyping PCR and ddPCR

Genomic DNA (gDNA) was extracted from hPSCs with the DNeasy Blood and Tissue kit according to manufacturer’s instructions (Qiagen). For genotyping of WT AAVS1 in dCas9-KRAB and dCas9-VPR clones, PCR of gDNA was performed with primer pair GE222 and GE668. For genotyping of gene targeted AAVS1 in dCas9-KRAB cells, PCR was performed with primer pair GE222 and GE586 for 5’ junctions and primer pair GE819 and GE668 for 3’ junctions. For genotyping gene targeted AAVS1 in dCas9-VPR cells, PCR was performed with primer pair GE222 and GE332 for 5’ junctions and primer pair GE233 and GE668 for 3’ junctions. For ddPCR of gDNA to quantify *piggyBac* copy number in dCas9-KRAB-PB and dCas9-VPR-PB clones, 20 μl reactions were prepared with ddPCR Supermix for Probes (no dUTP) (Bio-Rad, #1863024) with probes specific to mRFP and control gene *RPP30* according to manufacturer’s instructions (Bio-Rad). Droplets were generated using a QX100 Droplet Generator and PCR was performed on a C1000 Touch thermal-cycler (Bio-Rad) followed by sample streaming onto a QX100 Droplet Reader (Bio-Rad). Quantification was performed with QuantaSoft software. Primer sequences are listed in Supplemental Table S1.

### RT-qPCR

Total RNA from hPSCs was extracted using an RNeasy Mini Kit (Qiagen). Reverse Transcription cDNA synthesis reactions were performed on 0.2 μg −2 μg total RNA with iScript cDNA synthesis kit (BioRad) according to manufacturer’s instructions. Quantitative PCR reactions were performed the iTaq Universal SYBR Green Supermix (BioRad) and quantified by the ΔΔcT method on a CFX384 Real-Time System (Bio-Rad). Primer sequences are listed in Supplemental Table S1.

### Embryoid body differentiation and immunostaining

Embryoid bodies (EBs) were generated as previously described (Hazelbaker et al., 2017). For immunostaining, hPSC colonies and EBs were fixed with 4% paraformaldehyde in PBS for 15 mins at room temperature (RT), blocked and permeabilized with 0.1% TritonX-100 and 4% serum in PBS for 1 hr at RT and incubated with the appropriate primary antibody at RT. Following primary antibody incubation, cells were washed with PBS and incubated with the appropriate secondary antibody (Alexa Fluor 488 or 594, 1:500, Invitrogen) for 1 hr. Cells were then washed with PBS and incubated with DAPI before imaging at 20X magnification. The following primary antibodies were used: OCT4 (R&D Systems AF1759; 1:250), SSEA-4 (SCBT SC21704; 1:250), TRA-1-60 (SCBT SC21705; 1:200), AFP (Sigma A8452; 1:250), SMA (Sigma A2547; 1:2000), β-III Tubulin (R&D Systems MAB1195; 1:3000).

## RESOURCE DISTRIBUTION

All plasmids generated in this study including all-in-one dCas9-KRAB and dCas9-VPR targeting plasmids and multiplexed PB gRNA delivery systems will be deposited in Addgene.org upon publication. Cell lines will be made available upon request with appropriate institution approvals and following WiCell requirements for cell line distribution.

## AUTHOR CONTRIBUTIONS

D.Z.H., K.E., and L.E.B conceived and designed the study. A.B., P.M., G.A., A.M., D.L., and D.Z.H. performed the experiments and data analysis. D.Z.H. and L.E.B wrote the manuscript with input from all coauthors.

## ACKNOWLEDGEMENTS

This project was funded by the Stanley Center for Psychiatric Research at the Broad Institute. We thank Robert Ihry, Katie Worringer, Ajamete Kaykas (Novartis), Jesse Engreitz (Broad Institute), and Alejandro Chavez (Columbia University) for sharing of plasmids and cloning suggestions. We thank Tõnis Timmusk (Tallinn University of Technology) for TCF4 isoform identification advice. We thank members of the Barrett laboratory for advice and suggestions. We thank the Broad Institute Flow Facility for experimental support.

---

**Figure S1.**
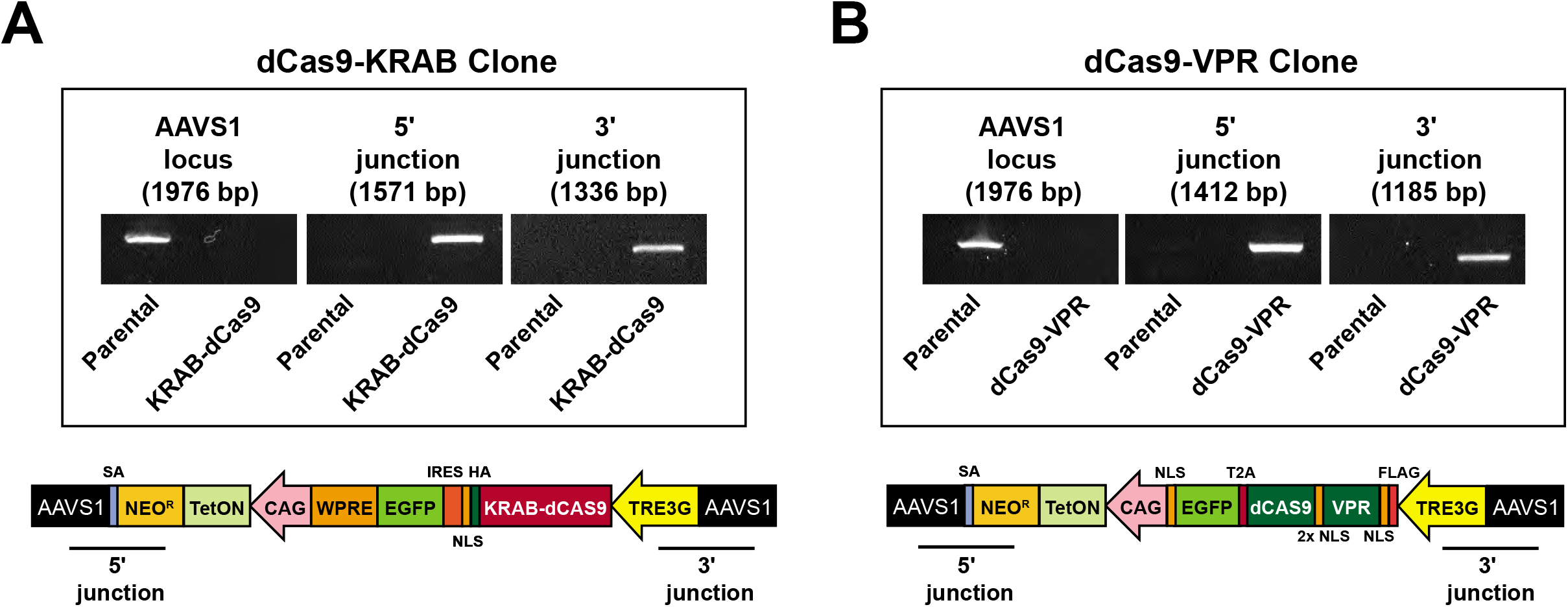
**A, B.** Genotyping of AAVS1 integration in dCas9-KRAB and dCas9-VPR parental clones by junction PCR.

**Figure S2.**
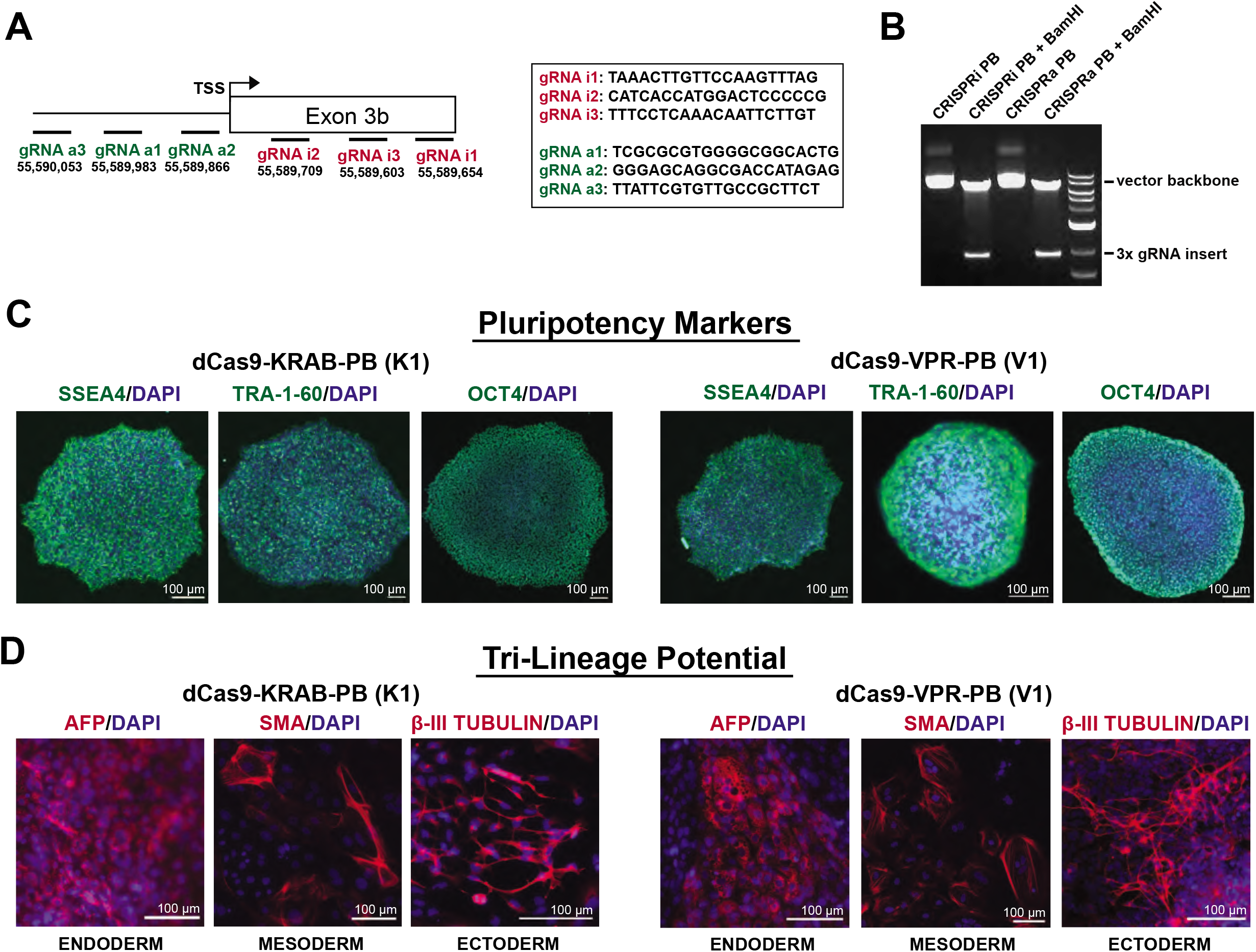
**A.** Relative locations and sequences of *TCF4* gRNAs. Numbers correspond to genetic coordinates in hg38 human genome assembly. **B.** Confirmation of presence of 3x gRNA insert in multi-gRNA PB vectors by digestion with BamHI restriction enzyme. **C.** Representative immunostaining for pluripotency markers SSEA4, TRA-1-60, and OCT4 in dCas9-KRAB-PB and dCas9-VPR-PB cells. **D.** Representative immunostaining for AFP (endoderm), SMA (mesoderm) and β-III Tubulin (ectoderm) following embryoid body formation from dCas9-KRAB-PB and dCas9-VPR-PB cells. Cells are counterstained with DAPI.

**SUPPLEMENTAL TABLE S1.**
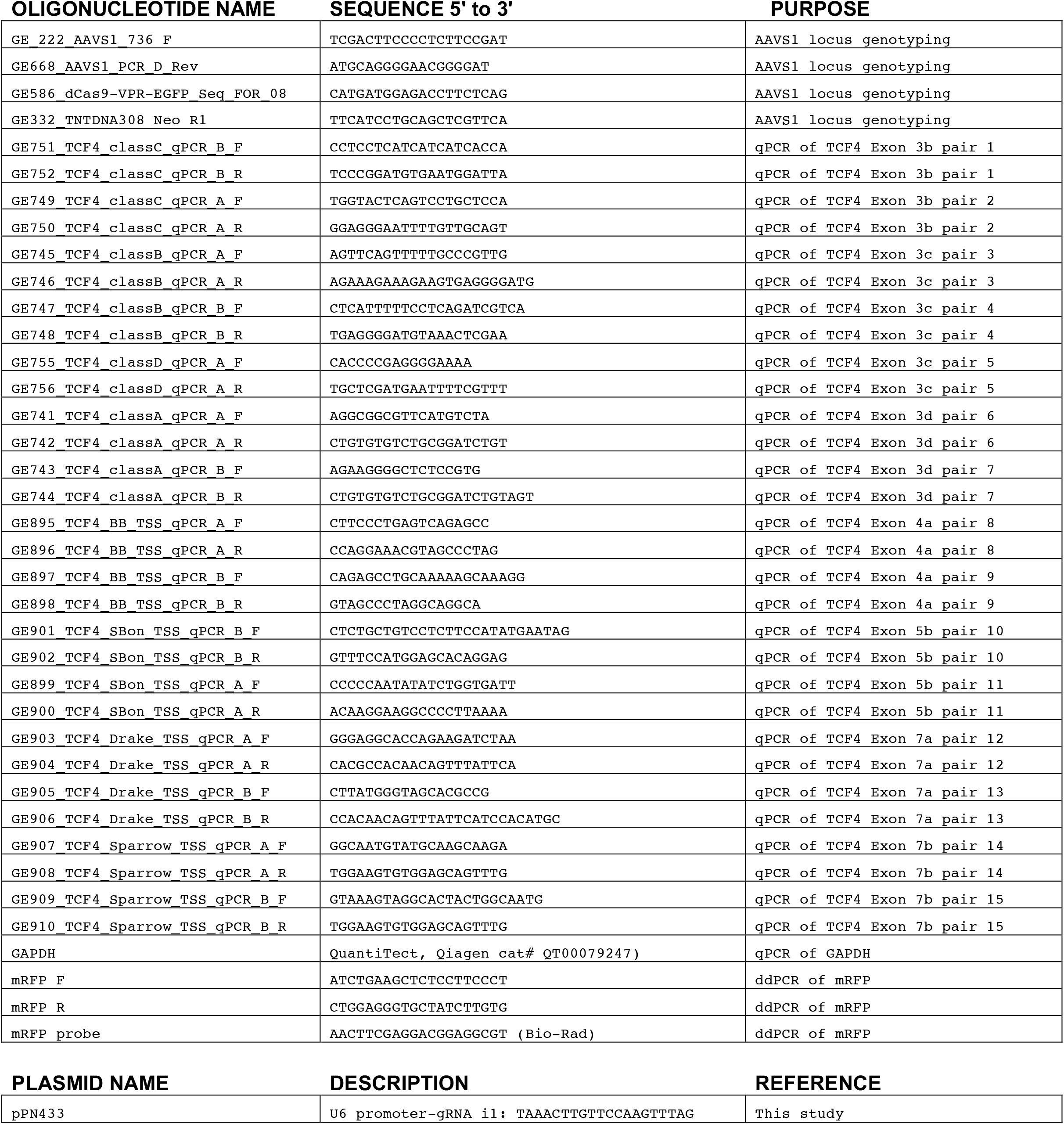
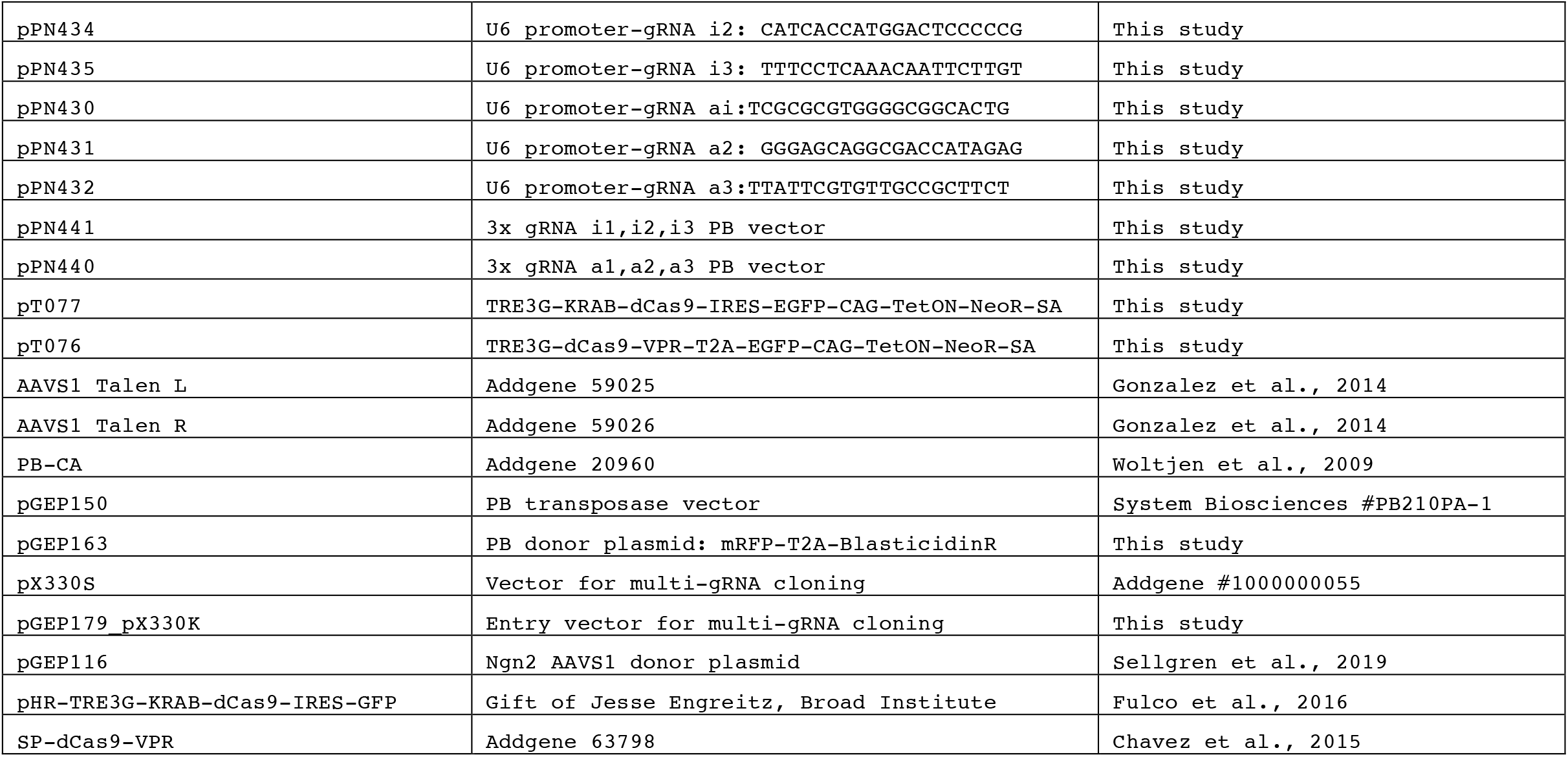
Oligonucleotides and plasmids used in study.

